# Local muscle pressure stimulates the principal receptors for proprioception

**DOI:** 10.1101/2024.02.06.579058

**Authors:** Frida Torell, Michael Dimitriou

## Abstract

Proprioception plays a crucial role in motor coordination and self-perception. Muscle spindles are the principal receptors for proprioception. They are believed to encode muscle stretch and signal limb position and velocity. Here, we applied percutaneous pressure to a small area of extensor muscles at the forearm while recording spindle afferent responses, skeletal muscle activity and hand kinematics. Three levels of sustained pressure were applied on the spindle-bearing muscle when the hand was relaxed and immobile (‘isometric’ condition) and when the participant’s hand moved rhythmically at the wrist. As hypothesized to occur due to compression of the spindle capsule, we show that muscle pressure is an ‘adequate’ stimulus for human spindles in isometric conditions, and that pressure enhances spindle responses during stretch. Interestingly, release of sustained pressure in isometric conditions lowered spindle firing below baseline rates. Our findings urge a re-evaluation of muscle proprioception in sensorimotor function and various neuromuscular pathologies.

**HIGHLIGHTS:**

1. Local muscle pressure is an adequate stimulus for spindles in isometric conditions
2. Release of pressure in isometric conditions lowers spindle firing below baseline rates
3. Local muscle pressure enhances spindle afferent responses during muscle stretch

## INTRODUCTION

It is well established that intramuscular fluid pressure (IMP) increases with contractile force and passive muscle elongation^1-3^. IMP has proven a better index of muscle force than muscle electrical activity (Electromyography: EMG), especially so in the case of eccentric contractions, muscle fatigue and recovery^4,5^. It is also known that single motor unit activity involves sharp increases in pressure within the motor unit territory, with no effect at more distant sites in the muscle, and that IMP influences the degree of actively produced muscle force^6^. It follows that access to sensory feedback concerning muscle pressure gradients would empower more robust sensorimotor function, such as by directly incorporating the muscle’s fluidic behavior in segmental reflex arcs and more global muscle activation strategies^7^. However, with regard to skeletal muscle proprioception, it is generally believed that the nervous system receives major sensory feedback only about stretch and global tension via muscle spindles and Golgi tendon organs, respectively (e.g., ^8,9^).

Muscle spindles are considered the principal receptors for proprioception^8,10^. Spindles can encode muscle stretch^11^ and their afferent signals have been implicated in a variety of sensorimotor functions including reflex control and the update of internal models of limb dynamics^12,13^. Recent studies have suggested additional functions for proprioceptive feedback, including in the maintenance of spinal alignment^14^ and the proper healing of fractured bones^15^. Moreover, in the early 1960’s, Bridgman and Eldred put forward a hypothesis that the secondary ending of the muscle spindle organ may be able to function as a pressure-sensing mechanism^16^. They argued that pressure acting on the expanded equatorial area of the spindle organ will compress and elongate the fluid-filled capsule, stretching the intrafusal fibers terminating inside the capsule (the more numerous “nuclear chain” fibers^17,18^), stimulating the spindle receptor ending. In fact, besides slow adaptation and sensitivity to passive stretch, response to well-localized press or ‘poking’ of muscle has long been the primary means of identifying muscle spindles in human and animal experiments (e.g., ^19-21^). However, there has been no systematic investigation of the ability of spindles to encode locally applied muscle pressure. Here, we assess an updated version of the original hypothesis: we test whether primary and secondary spindle receptors of awake humans can encode levels of muscle pressure (i.e., both endings engage nuclear chain fibers^22^), and also assess the impact of pressure on spindle responses during stretch.

Specifically, in this study we applied perpendicular, percutaneous pressure on one of three major extensor muscles of the forearm using a hand-held probe while recoding spindle afferent activity from this muscle (i.e., in the right radial nerve), as well as monitoring EMG and hand kinematics (Figure 1A). Three target levels of sustained press force were sequentially applied when the hand was relaxed and immobile (‘isometric’ condition) and when the participant’s hand was moved rhythmically about the wrist by the experimenter in the flexion-extension dimension at a 0.5 Hz pace (‘sinusoidal’ condition; Figure 1B). Documenting spindle responses to pressure when the hand was unmoving and during sinusoidal movement allowed us to study the impact of pressure in isometric conditions and in the context of muscle stretch during continuous movement. To confirm that any fusimotor control of spindles linked to action did not alter the basic impact of pressure during continuous movement, we also present three spindle afferent recordings where participants performed the sinusoidal movements actively.

**Figure 1.**
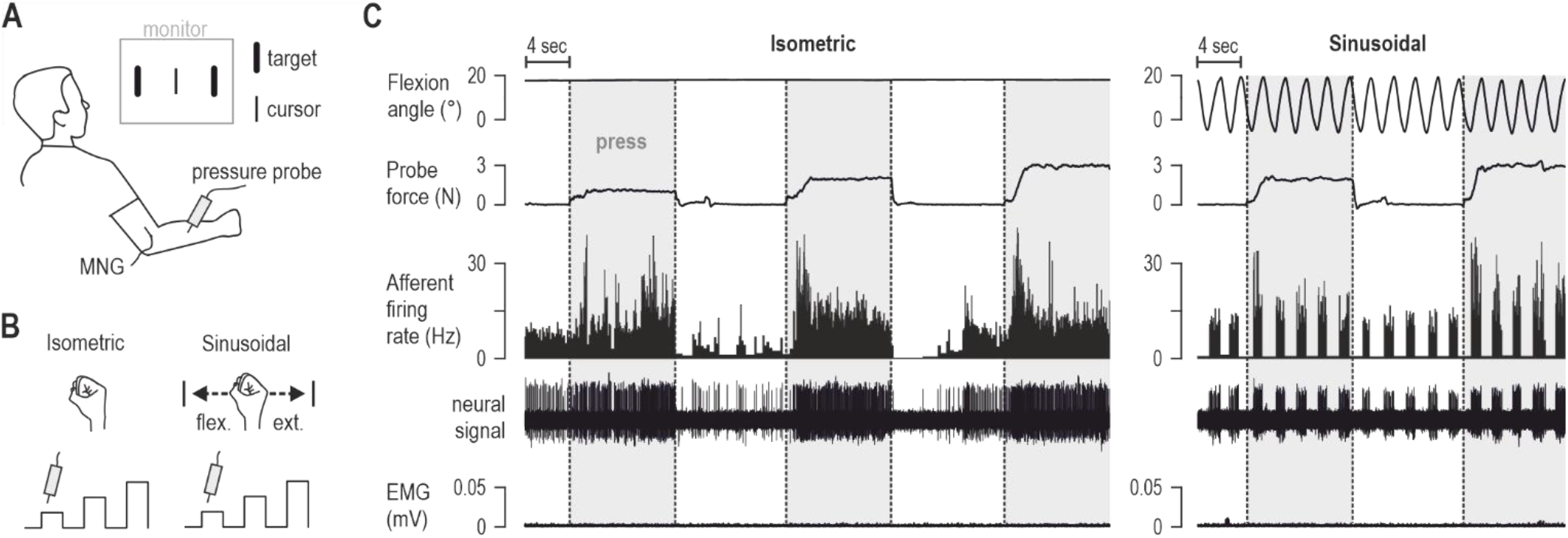
Experimental set-up and exemplary spindle afferent responses. **A)** Microneurography (MNG) was used for recording from muscle spindle afferents (n=14) in the right radial nerve of 8 awake individuals, while a probe applied pressure on a small part of the forearm (circular tip, ∼5mm diameter; see Methods section for more details). A kinematic sensor attached to the dorsum of the participant’s hand tracked movement about the wrist in the flexion-extension dimension. Surface EMG electrodes recorded muscle activity of wrist and finger extensor muscles from which the recorded spindle afferents originated. A monitor displayed two immobile visual targets and a cursor that tracked hand movement. **B)** The experimental task involved two conditions. In the first condition (‘isometric’), the semi-pronated hand was immobile whereas in the ‘sinusoidal’ condition the experimenter moved the participant’s hand continuously in the flexion-extension dimension so that the cursor reached the visual targets at a 0.5 Hz pace. In both conditions, three target levels of press force (1 - 3N) were sequentially applied, at the location on the forearm where the spindle afferent was most responsive to brief taps. Three of the participants also performed sinusoidal movements actively, for comparison of spindle behavior. **C)** Left: raw data of a spindle afferent from the extensor carpi radialis muscle during the isometric condition. Right: data from the same afferent/participant during the latter parts of the sinusoidal condition.

## RESULTS

The left panel of Figure 1C displays the responses of an exemplary spindle afferent recorded during the isometric condition. There is vigorous afferent firing in response to locally applied pressure on the spindle-bearing muscle. The right panel of Figure 1C shows that the same afferent responds with higher firing rates during wrist flexion (i.e., muscle stretch) when pressure is applied.

### The impact of pressure on spindle responses in isometric conditions

Figure 2A-C displays the responses of three additional muscle spindle afferents during the isometric condition, with each afferent originating from a different muscle (i.e., from the extensor digitorum, extensor carpi radialis and extensor carpi ulnaris, respectively). All three afferents respond to the locally applied pressure stimulus, with a dynamic component in their responses, and fall silent when sustained pressure is released in the ‘R2’ epoch.

**Figure 2.**
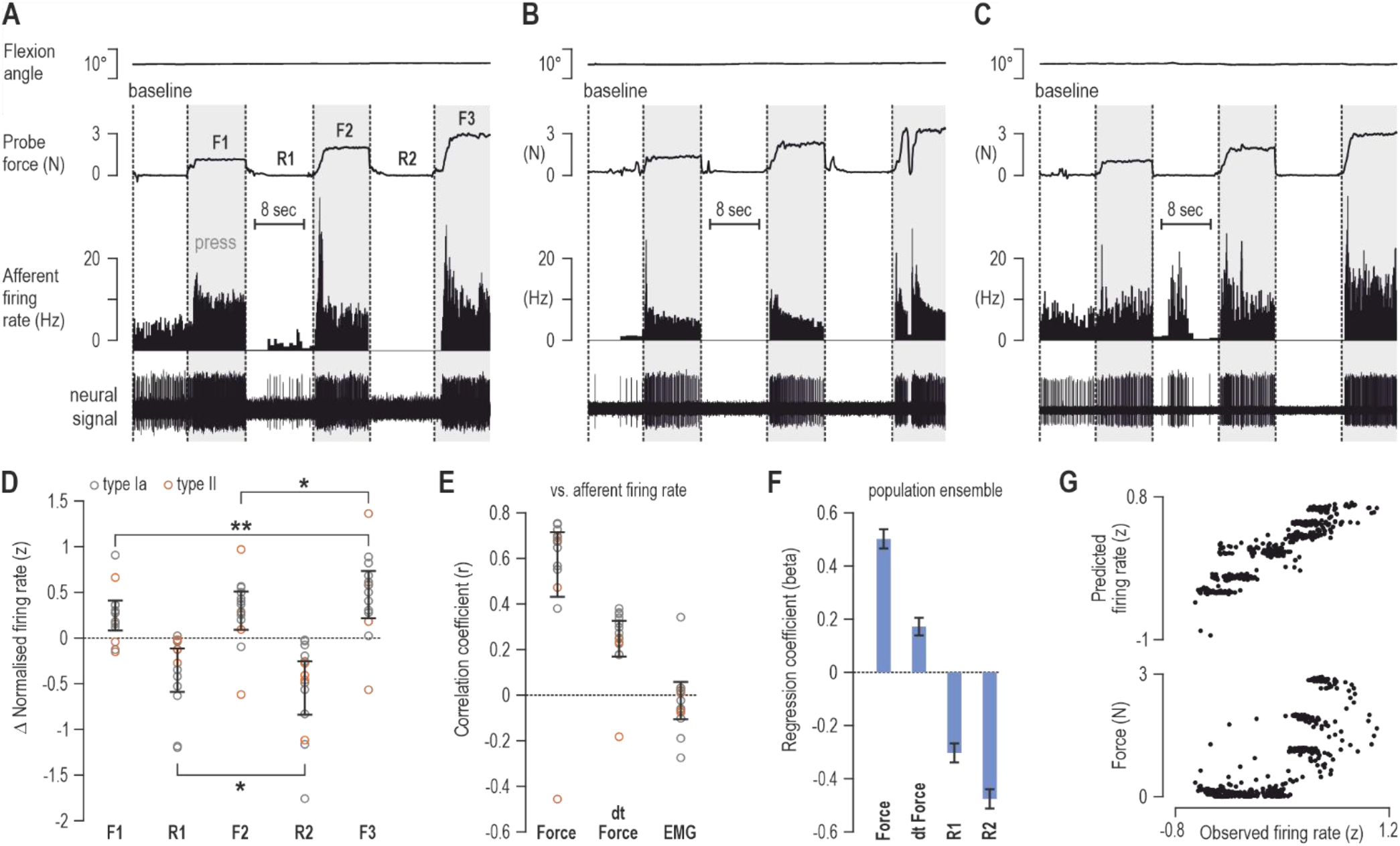
Local muscle pressure is an adequate stimulus for spindle receptors in the isometric condition. **A)** A spindle afferent from the common digit extensor muscle displaying vigorous responses to locally applied muscle pressure. For this analysis, the latter 8 seconds before application of the first press stimulus was designated as the baseline epoch, and the latter 8 seconds of the subsequent press and release epochs were coded as illustrated (i.e., F1-F3). **B)** As ‘A’ but representing a spindle afferent from the extensor carpi radialis muscle. **C)** As ‘A’ but representing a spindle afferent from the extensor carpi ulnaris muscle. **D)** Mean change in afferent firing rate (from baseline) in the five epochs of the isometric condition, as designated in ‘A’. Each circle outline corresponds to a single afferent (N=14). Error bars represent 95% confidence intervals. Statistically significant differences between individual groups are indicated by the lack of overlap between error bars (i.e., all p<0.01) or by asterisks in the case of overlapping bars. Single asterisk indicates statistical significance at alpha=0.05 and double asterisk significance at alpha=0.01. **E)** Full data sets during the isometric condition (e.g., see ‘A’-’C’) were down-sampled with a moving average window of 100 ms to perform correlations at the level of individual afferents. Error bars represent 95% confidence intervals, that were Fisher z transformed and then reversed to r. Only one afferent displayed a negative relationship between its firing rate and sustained press force or the first derivative of force (‘dt force’; see also Figure S2). **F)** Statistically significant beta (scaleless) regression coefficients following a forward stepwise regression to predict afferent population ensemble responses. All recorded afferents were used to form the ensemble, including the one represented in Figure S2. Error bars represent 95% confidence intervals. **G)** Above: the regression coefficients in ‘F’ produced a very good reconstruction of the observed ensemble firing rate. Below: a significant relationship between ensemble firing rate and the ensemble press force signal.

Figure 2D shows the mean change in firing rate across all recorded afferents for the different epochs of the isometric condition (latter 8 seconds of each epoch; see also Figure 2A). Data for this comparison were generated by first normalizing spindle firing rate (z-transform), and then subtracting the mean firing rate observed during the epoch before any pressure was applied i.e., ‘baseline’ epoch, at the level of single afferents. The recorded afferents were classified as primary (‘type Ia’) or secondary (‘type II’; see Methods for more details), although no systematic differences as a function of pressure were subsequently observed between the afferent types. Single-sample t-tests indicated that spindle firing rate was significantly different from baseline during all subsequent epochs, with sustained local press inducing stronger responses (F1: *t*_13_=3.24, *p*=0.0064; F2: *t*_13_=3.08, *p*=0.0086; F3: *t*_13_=3.97, *p*=0.0016) and ‘release’ epochs associated with lower firing rates than baseline (R1: *t*_13_=-3.19, *p*=0.007; R2: *t*_13_=-4.03, *p*=0.0014). A repeated-measures ANOVA of the three press epochs revealed a significant impact of pressure level (F_2,26_=7.4, *p*=0.003), with Tukey’s HSD test indicating significantly higher firing rates at the F3 epoch (vs. F1: *p*=0.003; vs. F2: *p*=0.022). Moreover, a dependent-samples t-test indicated significantly lower firing rates in the R2 epoch vs. R1 (*t*_13_=2.27, *p*=0.04). To statistically confirm the spindle’s dynamic response to local pressure (i.e., higher firing rates during the rising phase of the pressure stimulus vs. hold phase; see e.g., Figures 2A-C), we calculated the dynamic index^23^ for each afferent and press epoch (see Methods for more details). Single-sample t-tests using the dynamic index data indicated significantly higher firing rates during the rising phase of the pressure stimulus vs. hold (F1: t_13_=5.16, p=0.0002; F2: t_13_=6.8, p<10^−4^; F3: t_13_=3.6, p=0.003; Figure S1A). In addition, a repeated-measures ANOVA of mean firing rates observed during the rising phase of pressure yielded a significant effect of pressure level (F_2,26_=4.2, p=0.027; Tukey’s HSD test indicating F3 > F1, p=0.023; Figure S1B). Overall, our analyses suggest that local pressure is an ‘adequate’ stimulus for muscle spindles in the isometric condition, with significantly higher firing rates for higher press forces (∼2N resolution in our study; but see also ‘Limitations’), and release of pressure is associated with lower firing rates than baseline.

To complement these findings, we also analyzed continuous raw data profiles of single afferents (see e.g., Figure 2A-C), down-sampled with a moving average window of 100 ms. As illustrated in Figure 2E, the firing rate of all but one afferent displayed a positive relationship with press force (95% CI in r [0.71, 0.43]) and with its first derivative (95% CI in r [0.33, 0.17]). Although participants were instructed to remain relaxed during the isometric condition, some minor EMG activity was occasionally evident in the spindle-bearing muscle of some participants. The relationship with EMG activity was therefore also examined as a proxy for alpha-gamma co-activation^24^. However, as expected, there was no systematic relationship between EMG and spindle firing across the recorded afferents (95% CI in r [0.06, -0.11]). The down-sampled profiles were also collapsed across all afferents (averaged) to create population ensemble signals for a forward-stepwise regression. Specifically, press force and its first derivative, as well as dummy variables representing the release epochs (‘R1’ and ‘R2’) and spindle-bearing muscle EMG were entered as predictors to reconstruct the afferent ensemble responses. A very good reconstruction of ensemble firing rate was achieved (adjusted r^2^=0.85, p<10^−5^), with all predictors except EMG exerting a significant impact (force β=0.5, p<10^−5^; force dt β=0.18, p<10^−5^; R1 β=-0.3, p<10^−5^; R2 β=-0.47, p<10^−5^; EMG β=0.02, p=0.19; Figure 2F-G). A strong positive relationship is also evident between the population firing rate and press force, with r=0.75 and p<10^−5^ (Figure 2G, bottom). Performing the above analyses using data down-sampled with a larger moving average window (500 ms) produced virtually identical results (Figure S3). Taken together, our results demonstrate that in isometric conditions spindle neurons are strongly affected by and can encode locally applied muscle pressure, whereas release of sustained pressure lowers spindle firing below baseline rates.

### The impact of muscle pressure on spindle responses during stretch

Figure 3A displays the responses of a typical spindle afferent recorded during the sinusoidal condition (see also Figure 1C, right). In this condition, there is no increase in background firing throughout periods of pressure; rather, the afferent responses to muscle stretch (wrist flexion) are stronger during periods of pressure. Figure 3B shows the mean change in afferent firing rates across all recorded afferents for the different epochs of the sinusoidal condition. Data for this comparison was generated as per Figure 2D. Single-sample t-tests confirmed that spindle firing rates were significantly higher than baseline during all epochs where pressure was applied (F1: *t*_13_=2.9, *p*=0.012; F2: *t*_13_=2.97, *p*=0.011; F3: *t*_13_=3.83, *p*=0.002). Moreover, a repeated-measures ANOVA of the three press epochs again produced a significant impact of pressure level (F_2,26_=6.3, *p*=0.006), with Tukey’s HSD test indicating significantly higher firing rates in the F3 epoch vs. F1, with *p*=0.004. In contrast to the isometric condition, however, single-sample t-tests indicated no systematic difference in firing rate during the release epochs (R1: *t*_13_=-0.45, *p*=0.66; R2: *t*_13_=-0.32, *p*=0.75). As expected, performing the same single-sample t-tests as for spindle firing using wrist angle, angular velocity and muscle EMG data indicated no significant difference from baseline values in any epoch (all p>0.05).

**Figure 3.**
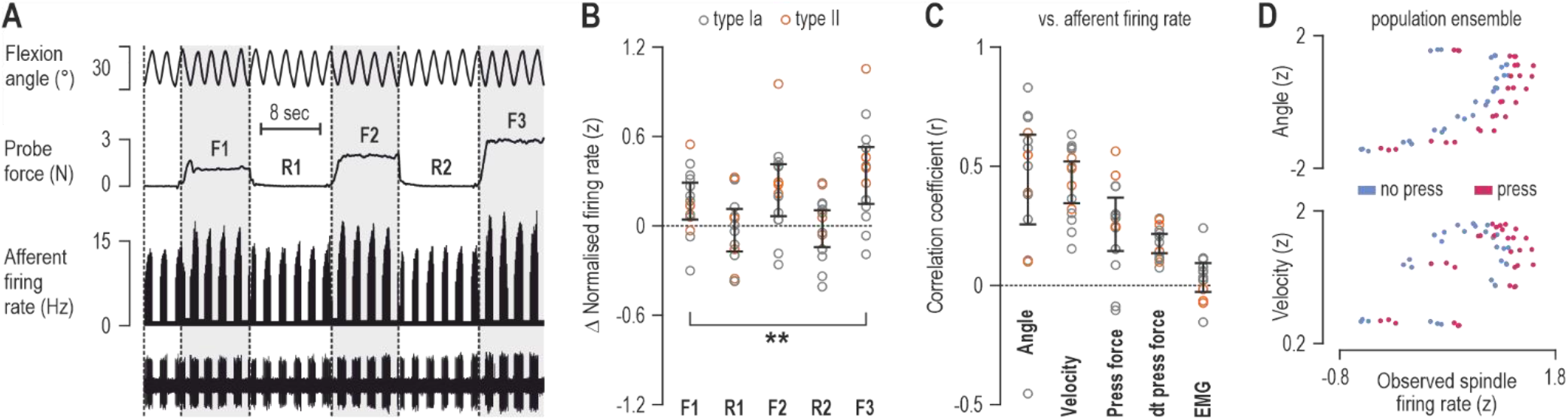
Local muscle pressure enhances spindle responses during stretch. **A)** Exemplary responses from a spindle afferent during the sinusoidal condition. Despite the same sinusoidal movement of the hand, higher firing rates are evident in response to muscle stretch (wrist flexion) when local press was applied on the muscle. As per the isometric condition, the latter 8 seconds before application of the first press stimulus was designated as the baseline epoch, and the latter 8 seconds of the subsequent press and release epochs were coded as illustrated (i.e., F1-F3). **B)** Mean change in afferent firing rate (from baseline) in the five epochs of the sinusoidal condition, as designated in ‘A’. Each circle outline corresponds to a single afferent (N=14). Error bars represent 95% confidence intervals. Statistically significant differences between groups that deviate from baseline are indicated by asterisks. Double asterisk represents statistical significance at alpha=0.01. **C)** Full data sets in the sinusoidal condition were down-sampled with a moving average window of 100 ms and correlations were performed at the level of individual afferents. Error bars represent 95% confidence intervals that were Fisher z transformed and then reversed to r. **D)** Normalized ensemble population signals during muscle stretch in the sinusoidal condition. Red dots correspond to datapoints during the three press epochs (F1-3) and the blue dots represent the equivalent for the three epochs with no press (i.e., baseline and R1-2). At the population ensemble level (and single afferent level as well; see e.g., ‘A’), the impact of muscle pressure on spindles manifested primarily as a positive offset in afferent responses.

We also analyzed continuous data profiles of single afferents (see e.g., Figure 3A), down-sampled with a moving average window of 100 ms. However, since it was clear from the raw data profiles that vigorous afferent responses were present only during muscle stretch, these specific analyses were confined to periods of wrist flexion. As indicated in Figure 3C, the firing of the majority of afferents displayed a positive relationship with flexion angle (95% CI in r [0.63, 0.26]), angular velocity (95% CI in r [0.52, 0.34]), press force (95% CI in r [0.37, 0.14]) and its first derivative (95% CI in r [0.22, 0.14]), but there was no systematic relationship between afferent firing and EMG (95% CI in r [0.1, -0.03]). Population ensemble signals pertaining to the variables named in Figure 3C were used to reconstruct the afferent population firing rate (see Methods for more details). Following a forward stepwise regression, a very good reconstruction of ensemble firing rates was achieved (adjusted r^2^=0.95, p<10^−5^) with joint angle, angular velocity and press force emerging as significant predictors (angle β=0.77, p<10^-5^; velocity β=0.7, p<10^−5^; force β=0.31, p<10^−5^). As shown in Figure 3D, the impact of pressure manifested as a positive offset in afferent responses during wrist flexion; there were otherwise no significant differences in r values between spindle firing rate and angle or velocity as a function of press i.e., angle red r (0.61) vs. angle blue r (0.7), p=0.52; velocity red r (0.63) vs. velocity blue r (0.55), p=0.65.

### Contrast of spindle responses across conditions

Comparing the r^2^ values obtained for the relationship between single spindle firing and press force in the isometric condition (Figure 2E) with those obtained for angle and velocity (Figure 3C) indicated a significantly stronger relationship for the former (Figure 4). These results further reinforce that local pressure should be considered an ‘adequate’ stimulus for spindles in isometric conditions. Specifically, dependent-samples t-tests indicated that press force in the isometric condition was more strongly related to spindle firing than joint angle (*t*_13_=2.38, *p*=0.033) or angular velocity (*t*_13_=3.64, *p*=0.003) during sinusoidal movement. Shapiro-Wilk tests indicated that the distribution of r^2^ values relating to press force did not depart from normality (W=0.91, p=0.16), nor did the r^2^ values relating to angle (W=0.96, p=0.81) and angular velocity (W=0.93, p=0.35). Last, we recorded from three spindle afferents during active sinusoidal movement, and local pressure again resulted in enhanced spindle responses during stretch (Figure S4).

**Figure 4.**
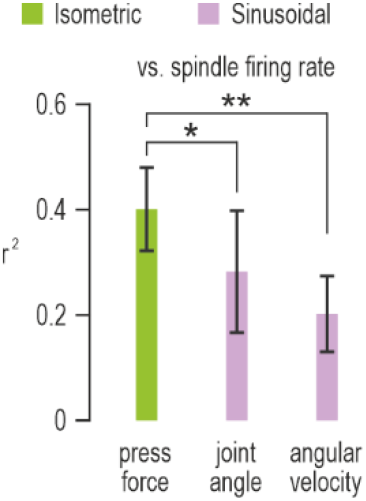
The closest observed relationship is between spindle firing and press force in the isometric condition. Mean r^2^ values across afferents (generated from the r values shown in Figure 2E and Figure 3C) describing the relationship between spindle afferent firing and press force in the isometric condition (green) and the relationship between spindle firing and joint angle/velocity in the sinusoidal condition (pink). Error bars represent 95% confidence intervals. Single star signifies statistical significance at alpha=0.05, and double star signifies statistical significance at alpha=0.01.

## DISCUSSION

This study demonstrates that muscle spindles can encode local muscle pressure in isometric conditions, and that release of sustained pressure in such conditions decreases spindle firing below baseline rates (Figure 2). We also show that pressure enhances spindle responses during muscle stretch (Figure 3). While we used percutaneous press to stimulate spindle receptors in our study with fully alert humans (see also ‘Limitations’), evidence in support of the hypothesis that compression of spindle capsules occurs during muscle contraction has been presented long ago^16^. IMP is naturally associated with both muscle contraction and percutaneous compression, such as when sitting or leaning on muscles in everyday life. Muscle spindles may routinely encode local press force and IMP in particular, with the latter known to be a more reliable index of muscle force than muscle electrical activity^4,5^. Spindle receptor endings are supplied with the stretch-sensitive Piezo2 channel^25^, which would argue for a predominant role in sensing muscle stretch. But Piezo2 is the main channel involved in peripheral mechanoreception^26,27^, including in skin receptor organs such as the Merkel disk receptor^28^ which is known to encode pressure and texture^8,29^. In other words, the structure and placement of encapsulated sensory organs can be decisive in terms of shaping their afferent signals.

Our findings demonstrate that spindles encode sustained pressure especially well during isometric conditions. Moreover, we found that the positive relationship between afferent firing and press force in the isometric condition is stronger compared to that between afferent firing and kinematics during muscle stretch (Figure 4). This reinforces the notion that local pressure should be considered an ‘adequate’ stimulus for muscle spindles. Spindle responses to press generally involved a positive offset in firing which could reflect mechanical stimulation of nuclear chain fibers^18,30^, as initially proposed by Bridgman and Eldred^16^ (see also Introduction section). However, subsequent work established the presence of three distinct types of intrafusal muscle fibers in mammalian spindles^17^. How the three fiber types individually contribute to the spindle’s response to pressure remains to be determined, and can be directly addressed using animal models (see ‘Limitations’). Moreover, the observed increase in absolute firing rates in response to pressure was variable across single afferents. This variability likely depends on multiple factors, such as the physiological characteristics of the particular spindle organ, muscle length and the depth of the spindle capsule beneath the muscle surface. One exceptional afferent responded negatively to sustained pressure; note, however, that even this single afferent responded positively during the rising phase of the pressure stimulus (Figure S2). We can only speculate as to the response pattern of the single exceptional afferent. One possibility is that local pressure was unsuccessfully applied in a sustained manner proximal to the spindle due to fascial interactions^31^. Nevertheless, in addition to the systematic positive effect of sustained press across the afferent population, there were even higher firing rates during the rising phase of the pressure stimulus (Figure S1A). That is, the spindle afferents also displayed a dynamic sensitivity to local pressure applied in isometric conditions. The dynamic component could possibly reflect stimulation beyond nuclear chain fibers e.g., some nuclear bag fibers also terminate near the polar ends of the spindle capsule^32^.

During continuous movement, the impact of pressure manifested as an increase in afferent firing during periods of muscle stretch alone. In other words, the known negative effect of muscle shortening on spindle responses seemed to override any positive effect of sustained pressure in this study. Fundamentally the same afferent response patterns were observed when continuous movement was executed actively (Figure S4). Spindles are unique peripheral sensory organs in that their output can be controlled independently by the nervous system itself, via gamma motor (fusimotor) neurons^33,34^. Over the years, several hypotheses have been put forward regarding the role of spindle control, including the maintenance of sensitivity to stretch^35^ via coactivation of fusimotor neurons^24^. But such presumed ‘alpha-gamma coactivation’ is able to maintain spindles responsive only when muscle shortening is slow^36^ or when the opposing load is large^37^. We did not vary the speed of movement in our study, but one prediction stemming from our findings is that local pressure may induce effects normally attributed to alpha-gamma coactivation. That is, local pressure may increase or maintain spindle responses during muscle shortening if movements are relatively slow or pressure on the spindle capsule is large. Moreover, because muscle contraction is accompanied by increases in IMP^1-3^, it is possible that increases in spindle firing observed during isometric contraction^38^ largely result from increased levels of IMP (i.e., compression of the spindle capsule). Future studies will contrast the effect of local pressure with that of voluntary isometric contraction; the difference in afferent response profiles between the two conditions may more effectively isolate the impact of any fusimotor coactivation during isometric contraction.

The ability of spindles to encode muscle pressure urges a re-evaluation of the role of proprioceptive muscle feedback in sensorimotor function^9,39,40^ and various neuromuscular conditions^41^, including the case of exercise-induced muscle cramps^42,43^. Afferent signaling of local muscle pressure may serve different functions across different contexts. Regarding sensorimotor function, in particular, this ability may underlie a more robust ‘gain-scaling’ mechanism for postural reflex control. That is, spindle responses typically increase during voluntary isometric contraction of the spindle-bearing muscle^38^. This increase is congruent with the known “automatic gain-scaling” of stretch reflexes, where reflex sensitivity or ‘gain’ is proportional to the background activation level of the muscle homonymous with the reflex^44,45^. Whereas ‘antagonistic muscle balance’^46^ shapes stretch reflex gains during ballistic movement^47^, gain-scaling in postural tasks is stronger overall and dominated by the state of the homonymous muscle^45,47^. We propose that gain-scaling during postural tasks is partially determined by the ability of spindles to encode muscle pressure. Because IMP is a better metric of the true tensile state of muscle than muscle electrical activity^4,5^, the spindle’s ability to encode local press can empower a more robust gain-scaling mechanism compared to one that only reflects the degree of neural activation (i.e., alpha-gamma coactivation) regardless of passive muscle tension, muscle fatigue etc. But outside the realm of purely postural tasks, the spindles’ mechanical gain-scaling property may potentially interfere with independent fusimotor control of spindles and stretch reflexes. For example, in the context of delayed reach, we have repeatedly shown that goal-directed tuning of stretch reflexes is attenuated by muscle (pre-)loading^48,49^, whereas muscle unloading (i.e., assistive loads) facilitates such tuning^50^. It should also be noted that prolonged high levels of IMP as in the case of compartment syndrome^51^, or interventions inducing transient hypoxia and nerve compression such as using a pressure cuff may attenuate sensory signals and reflex responses^52,53^.

Our study also shows that release of pressure sustained over seconds under isometric conditions led to the decrease in spindle firing below baseline rates (i.e., lower mean firing rates than those observed before any percutaneous pressure was applied; see e.g. Figure 2A-C). This spindle behavior could be due to an alleviation or redistribution of pre-existing levels of IMP close to the spindle capsule and/or due to changes in the elements of the spindle organ itself. We can only speculate as to the underlying mechanisms giving rise to this inhibition. Nevertheless, given the spindles’ established role in regulating muscle tone and stiffness, our results indicate why a ‘triple-8’ technique (i.e., 8 sec 1N press - 8 sec release - 8 sec 2N press) followed by slow passive stretch could be effective in quickly reducing pain and stiffness associated with local ‘trigger and tender points’^54^ in skeletal muscle.

### Limitations

While our minimally invasive *in vivo* study allowed us to systematically assess the impact of local muscle pressure on spindle responses, there are limitations. In the current study, the force associated with percutaneous press was used as an approximation of the pressure applied on the spindle capsule inside human muscle. More invasive animal studies can elucidate further the impact of IMP on spindle compression and afferent signals from healthy and diseased muscle. For example, this can be achieved by using catheters to monitor local IMP during simultaneous recording of afferent feedback or via direct intramuscular compression of spindle capsules in more reduced preparations. Despite the approximate measure of spindle pressure used in the current study, we obtained quite strong correlations overall between single afferent firing and press force (Figure 2E); observing an even finer relationship between spindle responses and local pressure in reduced preparations is likely.

In the current study we opted for not randomizing the application order of press force magnitude. This increased the probability of securing spindle responses with at least one common experimental manipulation before each afferent recording was lost. The lack of stimulus randomization was not considered problematic in terms of convoluting the effect of pressure magnitude, because it has been previously shown that repeated application of the same local press stimulus gives rise to very similar spindle afferent responses^21^. Also note, in addition to analyzing categorical effects, we also analyzed continuous variables and consistently found close positive correlations between press force magnitude and single afferent responses. However, the current study cannot distinguish whether the progressive, below-baseline decrease in spindle firing observed when pressure was released (i.e., ‘R2’ vs. ‘R1’ epoch; Figure 2D) was due to the preceding press magnitude or due to the repeated application of press. A study where pressure magnitude is randomized can address this issue. Last, we used ramp-and-hold pressure stimuli with a relatively long hold period, to primarily investigate the impact of sustained pressure. But our recordings suggest that spindles can be most responsive to dynamic changes in pressure. Future studies will elucidate the ability of spindles to encode dynamic changes in local press e.g., by applying sinusoidal pressure stimuli in isolation, as well as in- and out-of-phase with sinusoidal stretch.

## ACKNOWLEDGMENTS

The authors would like to thank Carola Hjältén and Anders Bäckström for their technical support. This work was supported by grants awarded to M.D. by Hjärnfonden (FO2022-0308) and the Swedish Research Council (2020-02140). The funders had no role in study design, data collection, analysis or manuscript preparation.

## Author contributions

Conceptualization, M.D.; Methodology, M.D.; Investigation, M.D. and F.T.; Formal Analysis, M.D. and F.T.; Visualization, M.D. and F.T.; Writing – Original Draft, M.D. and F.T.; Writing – Review & Editing, M.D. and F.T.; Supervision, M.D.; Funding Acquisition, M.D.

## Declaration of interests

The authors declare no competing interests.

## Inclusion and diversity statement

We support inclusive, diverse, and equitable conduct in research.

## STAR METHODS

| REAGENT or RESOURCE | SOURCE | IDENTIFIER |
| --- | --- | --- |
| **Deposited data** |  |  |
| Analyzed data | This paper; Mendeley Data | DOI: 10.17632/549wt8drc7.1 |
| **Software and algorithms** |  |  |
| MATLAB | MathWorks, Natick | https://se.mathworks.com/products/matlab.html |
| Statistica | StatSoft Inc | https://www.statistica.com/en/ |

## RESOURCE AVAILABILITY

### Lead contact

Reasonable requests for further information and data resources should be directed to and will be fulfilled by the lead contact, Michael Dimitriou.

### Data and code availability

- Analyzed data sets have been deposited at Mendeley and are publicly available as of the date of publication. Accession numbers are listed in the key resources table. Multiple raw data profiles are displayed in graphical form throughout the paper (e.g., Figure 1-3; S2; S4).
- Any additional information required to analyze the data reported in this paper is available from the lead contact upon reasonable request.

### PARTICIPANT DETAILS

Eight neurologically healthy adult human volunteers participated in the current study (mean age 24.5 years ± 4.7 standard deviations; half self-identified as female and half as male). The participants provided written informed consent prior to participation, in accordance with the Declaration of Helsinki, and were financially compensated. No statistical method was used to determine sample size, but the number of participants (and single afferents) in the current study is similar or exceeds the number used in equivalent studies (see e.g.,^48,55^). Our study was conducted as part of a research programme approved by the Umeå Regional Ethics Committee.

## METHOD DETAILS

### Muscle afferent recordings

Microneurography (MNG) was used to record single spindle afferent activity in the radial nerve of the right arm^56,57,^, as previously done in our lab^46,48,58^. Afferent recordings were obtained from either the radial wrist extensor (extensor carpi radialis), the ulna wrist extensor (extensor carpi ulnaris) or the common digit extensor (extensor digitorum communis). The recorded action potentials were identified as originating from primary or secondary muscle spindles (vs. e.g., Golgi tendon organ or skin receptors) according to standard procedures described in detail elsewhere^59-61^. In short, these procedures include examining the afferent responses to ramp and hold stretches of the relaxed spindle-bearing muscle, the response to taps on the muscle (vs. skin of the hand) and assessing the variability in background firing rate^62^. In total, 14 muscle spindle afferents were recorded (10 ‘type Ia’ and 4 ‘type II’). Each participant contributed with at least one afferent recording. The number of afferents recorded here is similar or exceeds the number of afferents recorded in previous MNG studies with human participants^48,58,59,63^.

### Electromyography (EMG)

Custom-made surface EMG electrodes, with a diameter of 2 mm and an internal distance of 12 mm were attached to three forearm muscles. Specifically, recordings were made from the extensor carpi ulnaris, extensor carpi radialis and extensor digitorum communis. Electrode placement was selected using a hand-held stimulator probe and isometric contraction/relaxation maneuvers. Prior to placement the EMG electrodes were coated with conductive gel, and then attached with double-sided tape. A ground electrode was placed on the lateral epicondyle of the humerus bone.

### Experimental set-up and design

Participants sat on an adjustable dental chair with their right arm resting on a padded cushion. The chair was tilted backward and adjusted to fit the participant’s stature and shape. The forearm was secured in place using a padded wooden vise proximal to the wrist, and this allowed the hand to move freely while the arm was stabilized to prevent displacement of the MNG electrode. Visual feedback was provided by a monitor placed opposite the participant at approximately eye-level (see Figure 1A). Two vertical targets (26.5 cm long and 0.8 cm wide, located 8.5 cm from the origin) were continuously displayed on the monitor, and hand position was represented by a vertical cursor/line (25.5 cm long and 0.3 cm wide). Hand kinematics were recorded using a FASTRAK® sensor attached to the dorsal surface of the hand with double-sided tape and fixated with surgical tape. Movement of the semi-pronated hand about the wrist in the flexion-extension dimension controlled the 1D cursor position along the horizontal axis (i.e., between the visual targets).

The current study involved two experimental conditions (Figure 1B). In the first condition (‘isometric’), the participant’s hand was relaxed and immobile, resting on a cushion in a lightly flexed position at the wrist. Note that the label ‘isometric’ refers here to the lack of any externally perceptible movement of the participant’s hand. Before the onset of the second ‘sinusoidal’ condition, the experimenter first grasped the lightly fisted hand of the participant and then moved the participant’s hand in the initial position. In the initial position, the participant’s hand (i.e., third metacarpal joint) was aligned with the longitudinal axis of the forearm (i.e., zero flexion/extension and ulna/radial deviation), and this corresponded to the cursor being in the center of the screen, between the two visual targets. The sinusoidal task then followed, which involved the experimenter moving the participant’s lightly fisted hand rhythmically about the wrist in the flexion-extension dimension at a 0.5 Hz pace. The speed of hand movements was guided by a metronome and movement amplitude was guided by the visual targets i.e., the experimenter moved the cursor so that it alternated in reaching each of the two targets. On three occasions, single afferent recordings were made while the participants performed the sinusoidal movements actively, also using the metronome and visual targets to control the pace and amplitude of their movements (Figure S4). Throughout, a rotation of one degree at the wrist corresponded to a 0.7 cm movement of the cursor along the horizontal axis of the monitor. Because the distance between the center of the screen and the center of a target was 8.5 cm, a wrist rotation of ∼12 degrees was required for reaching the middle of each target from the initial position.

During each experimental condition, we applied pressure percutaneously on one of three major extensor muscles of the forearm using a hand-held probe (i.e., on the muscle from which the recorded single afferent originated). Each single afferent was recorded individually, at a different time, and pressure was applied only on the well-localized small area of the spindle-bearing muscle, over which the single afferent was most responsive to this mechanical stimulus. As the experimental design of this study was within-measures, both ‘isometric’ and ‘sinusoidal’ conditions were sequentially applied once for each recorded single afferent. Specifically, the press force on the muscle was applied using a probe with an embedded six-axis force/torque transducer (Nano F/T transducer; ATI Industrial Automation), and a flat circular contact area of 18 mm^2^ (4.8 mm diameter). As indicated by the schematic at the bottom of Figure 1B, after an idle period of 10 seconds (corresponding to the ‘baseline’ epoch; see e.g., Figure 2A) three sequential target levels of press force (1, 2, and 3N) were applied in each experimental condition. Press forces are reported throughout in terms of the perpendicular force applied via the hand-held probe. The onset and offset time for applying a pressure stimulus, as well as the magnitude of the stimulus for any press, was indicated to a second experimenter (other than the one moving the hand in the sinusoidal condition) by a message shown on a second monitor. The second monitor was placed across the second experimenter, and outside the visual field of the participant. Each ‘press’ message on the second monitor lasted for 10 seconds, as did the ‘release’ messages. Some minor discrepancy was to be expected between the duration of the visual messages and that of the actual press epochs, due to the reaction time of the second experimenter. However, in addition to performing complementary analyses with continuous data profiles (e.g., Figure 2E-G), analyses were confined to the latter 8 seconds of each epoch where necessary (e.g., Figure 2D), as the experimenter’s reaction time never exceeded 2 seconds. Concerning the magnitude of the applied press force, the experimenter received continuous real-time feedback in the form of a horizontal flat bar moving vertically along a background scale shown on the second monitor, with increases in force corresponding to movement of the bar towards the top of the scale.

Before inserting the MNG electrode, participants were familiarized with the two experimental conditions, including experiencing a dry run of the sinusoidal condition and performing the sinusoidal movement themselves. The participants were asked to remain relaxed and passive during the isometric and sinusoidal conditions. On three occasions where the afferent recording seemed particularly stable as the end of the passive sinusoidal movement (hence likely not to be lost during active movement), the participants performed the sinusoidal movement actively for comparison (Figure S4).

## QUANTIFICATION AND STATISTICAL ANALYSIS

### Data sampling and pre-processing

Data sampling was performed using SC/ZOOM™. With SC/ZOOM^™^ single action potentials were identified semi-automatically under visual control. The simultaneously recorded EMG channels were high-pass filtered with a 30 Hz cutoff and then root-mean-square processed with a symmetrical +-3.0 ms window. They were digitally sampled at 4800 Hz. All data used for analyses, including EMG, were extracted from SC/ZOOM^™^ at a 1000 Hz rate. To facilitate comparison of EMG and afferent recordings across muscles and participants, the raw data were normalized by z-transformation using a procedure that has been described elsewhere^58,59,64^. In brief, for each individual afferent/participant, and each data variable, a grand mean and grand standard deviation was generated across all continuous datapoints. The z-transformed data was obtained by subtracting the grand mean from the measured value and dividing the difference by the grand standard deviation. To generate the down-sampled data that was entered into correlations and regression analyses, a moving average window of 100 ms was used on the recorded raw data profile of each afferent (e.g., Figure 2A-C & 3A). To generate the population ensemble signals of the isometric condition, the down-sampled profiles were collapsed (averaged) across afferents (N=14). All afferents were used in this procedure, including the one with a net negative relationship to muscle pressure (Figure S2). Note that in one afferent profile (out of 14) the EMG signals alone were lost due to an acute battery issue with the recording equipment and therefore treated as missing data. Using a larger down-sampling window produced equivalent correlation and regression results (Figure S3). To generate the population ensemble signals pertaining to the sinusoidal condition (i.e., Figure 3D), a different approach was taken. For each single afferent separately, the raw data for each movement cycle was identified and re-sampled to 2 sec (corresponding to the target pace of movement: 0.5 Hz), and then collapsed (averaged) across the number of repetitions for each epoch (i.e., 6 epochs: baseline and F1-F3 as designated in Figure 3A). The data for each of the 6 averaged cycles per afferent where then down-sampled with a moving average window of 100ms (resulting in 20 data points per cycle), and finally averaged data cycles were collapsed across afferents to create the population ensemble signals, totaling 120 data points for each variable (6 epochs x 20 points each cycle; Figure 3D). Throughout, data pre-processing was performed using Matlab® (R2019a, MathWorks, Natick, MA, USA).

### Statistical analyses

To assess the impact of press force on spindle afferent firing, all statistical analyses were two-tailed, where applicable. These involved single-sample t-tests to assess significant deviation in firing rate from baseline levels, as well as repeated-measures ANOVA to assess the effect of pressure level, along with Tukey’s posthoc analyses and dependent-samples t-tests to compare firing rates across epochs (Figure 2D & 3B). Single-sample t-tests were also employed to confirm the lack of significant deviation in wrist angle, velocity and EMG from baseline as a function of task epoch, as well as for analyzing a dynamic index of each afferent (Figure S1A). The dynamic index^23^ is a widely used metric for assessing the responsiveness of afferents to dynamic changes in stimulation. As commonly generated, the dynamic index for each afferent and press epoch in this study was calculated as the difference in mean firing rate (z) observed during the rising phase of the stimulus (“ramp”) and the mean firing rate observed for an equal-sized period immediately following the end of the ramp phase. In cases where two rather distinct rise phases were evident (i.e., initial ramp was incomplete), the latter part of the ramp was used that subsequently led to a hold phase. For each afferent, the onset and termination of ramp phases were identified semiautomatically under visual control. Pearson’s correlation coefficients (‘r’) were calculated to assess the relationship between firing rate and various independent variables, both at the single afferent level and at the population level (Figure 2E,G & 3C). Dependent sample t-tests were used to contrast the strength of relationship between firing rate and press force in the isometric condition and that between firing rate and kinematic variables in the sinusoidal condition (Figure 4). Forward-stepwise regressions were used for identifying the significant predictors of afferent ensemble firing rate, using z-transformed data (Figure 2G & 3D). Specifically, regarding the isometric condition, the purpose of the complementary regression using population ensemble data was to assess the extent to which the variables with a consistent impact at the level of single afferents (i.e., press force and pressure release epochs) could together account for the observed variance in spindle population responses. Regarding the sinusoidal condition, press force and the kinematic variables commonly associated with spindle responses were together used for the same purpose. Beta coefficients are reported which represent the standard deviation change in the dependent variable (here afferent firing rate) for 1 standard deviation change in the predictor variable. To assess data normality, Shapiro-Wilks test was used for sample sizes <50 data points, otherwise the Lilliefors test was used. Throughout, the level of statistical significance was 0.05. Correlation analyses at the level of single afferents were performed using Matlab® (R2019a, MathWorks, Natick, MA, USA), whereas all other statistical analyses were performed using STATISTICA® (StatSoft Inc, USA).

## SUPPLEMENTAL FIGURES

**Figure S1.**
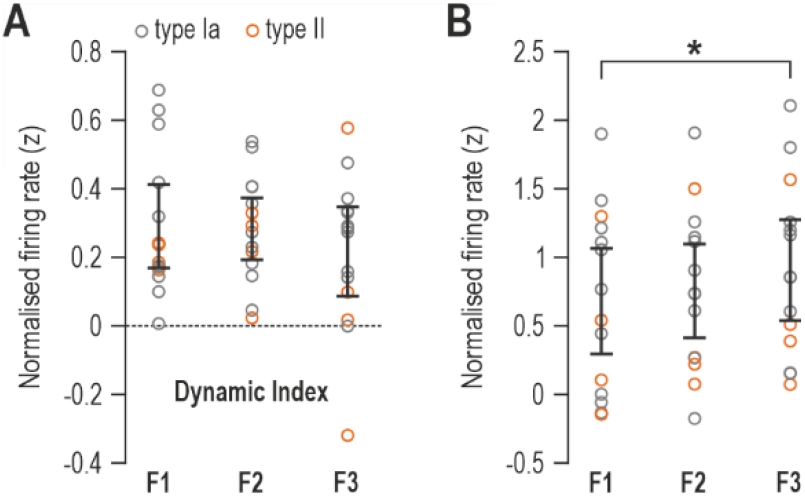
Single afferent responsiveness to dynamic increases in muscle pressure. **A)** The ‘dynamic index’ calculated for each afferent and press force epoch (F1-F3), indicating the relative sensitivity of spindles to the rising phase of the pressure stimulus (“ramp”) vs. the hold phase. Positive values indicate dynamic sensitivity (see Results and Methods for more details). Throughout Figure S1, error bars represent 95% confidence intervals. **B)** Mean afferent firing rates (z) during the ramp phase of pressure in the three press force epochs. The asterisk indicates a statistically significant difference at alpha=0.05.

**Figure S2.**
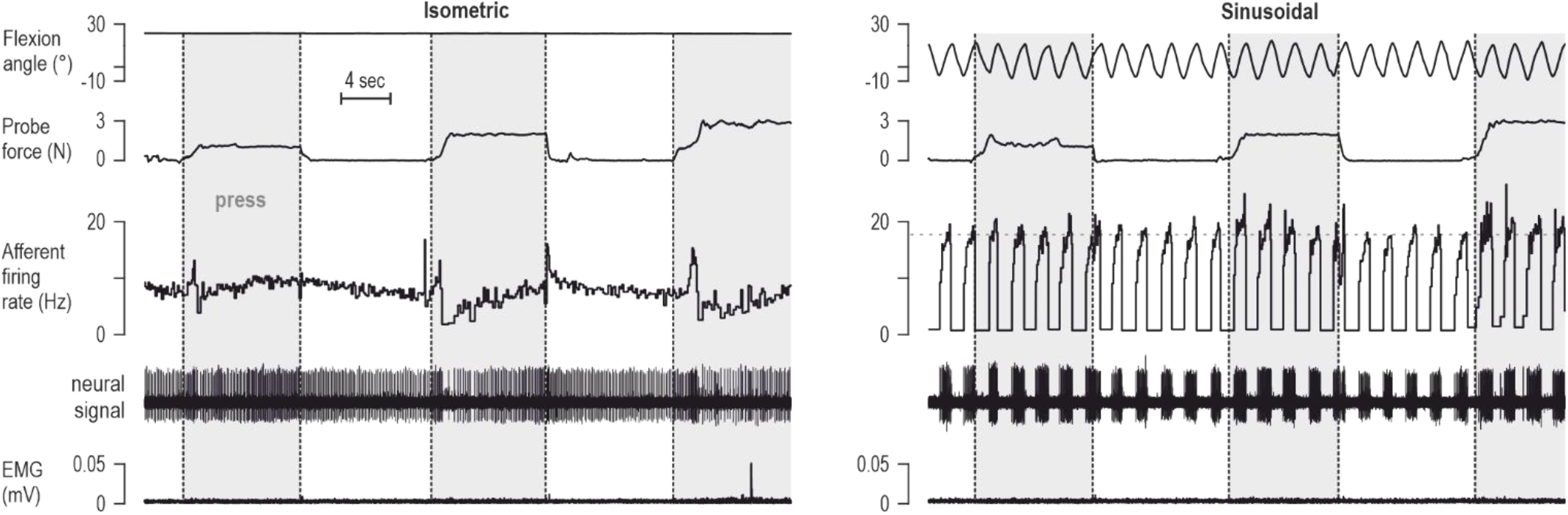
The only spindle afferent with a net negative response to muscle pressure in the isometric condition. Left: responses of the only recorded spindle afferent that displayed a negative overall relationship with press force in the isometric condition. Note, however, the brief increase in firing rate at the onset of each of the three press stimuli, indicating some dynamic sensitivity to pressure (i.e., the afferent did respond to brief taps at the location of the muscle where the probe subsequently pressed). Right: during the sinusoidal condition, the same afferent responded consistently to wrist flexion (i.e., stretch) with some minor increase in firing rate during the early movement cycles when the 2N and 3N press were applied.

**Figure S3.**
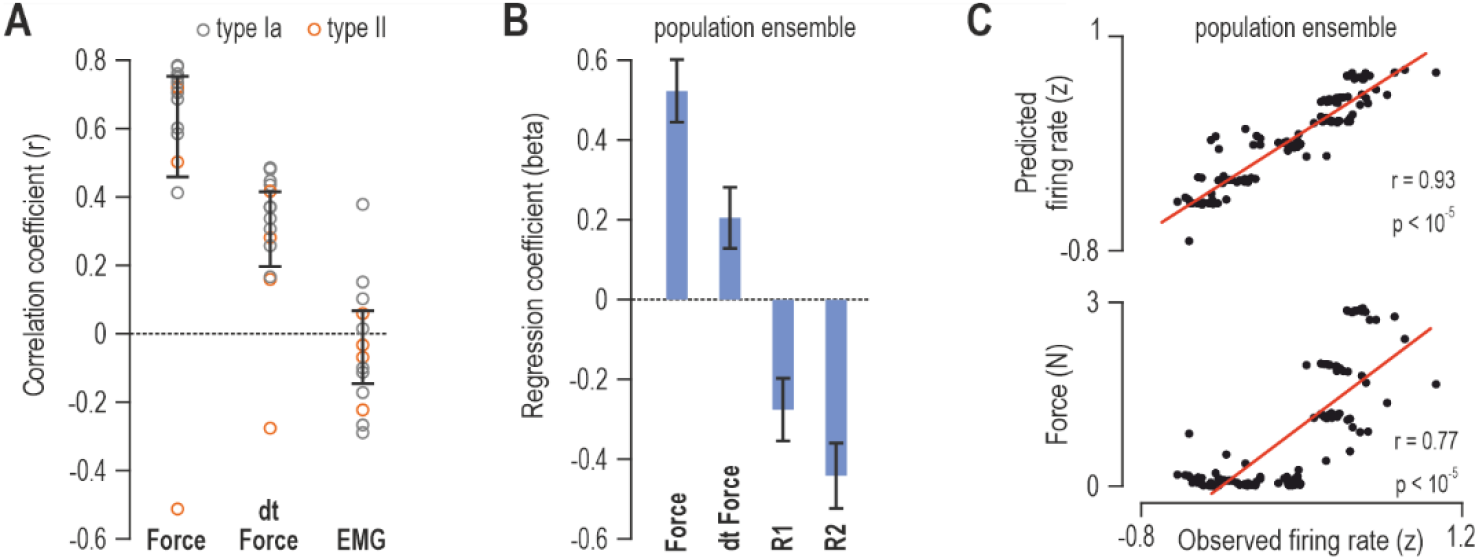
A larger down-sampling window produces qualitatively the same results. **A-C)** As Figure 2E-F but using a larger moving-average time window (500 ms) to down-sample the continuous data.

**Figure S4.**
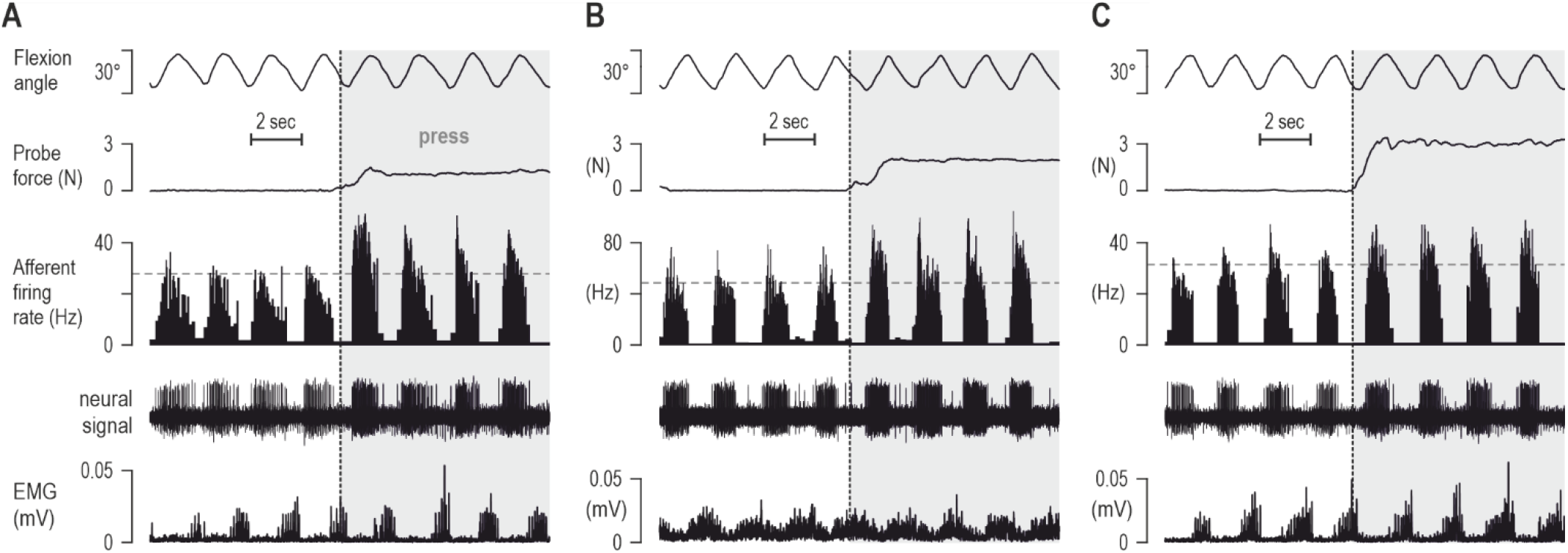
Local muscle pressure enhances spindle responses to stretch during active movement. **A-C)** Responses from three separate spindle afferents during active movement of the hand, as per the sinusoidal condition (e.g., Figure 3). Despite very similar movement of the hand in each case, higher firing rates are evident in response to muscle stretch (wrist flexion) when local press is applied on the muscle, especially for the afferents in ‘A’ and ‘B’.

## Notes

### Competing Interest Statement

The authors have declared no competing interest.

### Summary of Updates

Additional analyses (e.g., Figure S1) and an expanded discussion section

